# Theta oscillations in the human hippocampus normalize the information content of episodic memory

**DOI:** 10.1101/2022.06.27.497705

**Authors:** D. Santos-Pata, R. Zucca, A. Fernandez Amil, A. Principe, C. Pérez-Enríquez, R. Rocamora, S. C. Kwok, P. Verschure

## Abstract

The principles governing the formation of episodic memories from the continuous stream of sensory stimuli are not fully understood. Theoretical models of the hippocampus propose that the representational format of episodic memories comprise oscillations in the theta frequency band (2-8 Hz) that set the time boundaries in which discrete events are bound encoded in the gamma frequency range (>30 Hz). We investigated this temporal segmentation and binding process by analyzing the intracranial EEG (iEEG) of surgically implanted epileptic patients performing a virtual-navigation task. We found a positive correlation between sensory information density encountered by the subject and hippocampal theta-frequency, suggesting that the human hippocampus normalizes the information content of episodic memories relative to the density of sensory information. This interpretation is further supported by the observation that as a marker of mnemonic encoding, i.e. the amount of persistent gamma events, directly correlates with sensory information density, gamma-frequency power and the phase relation between theta and gamma oscillations remain constant. Using a theoretical model of the hippocampus, we build a model that analogously displays a similar normalization of gamma episodes per theta cycle relative to information density by accounting for the physiological signatures of theta-gamma coding through combining fast and slow inhibitory feedback. We propose that this intrinsic normalization mechanism optimizes the trade-off between the discretization and compression of continuous experience relative to the limited capacity of episodic memory.

**Summary:** We move in continuous time and space, yet we can encode and recall discrete episodes from our past experiences. The neural mechanism behind this discretization is not fully understood. It has been previously observed that rodent locomotion modulates ongoing hippocampal theta rhythms. Thus, raising the question of whether these slow rhythms bind events together during a single oscillatory cycle relative to the movement speed or overall information density.

We quantified the effects of increasing locomotion and sensory information in modulating theta oscillations during virtual navigation with intracranial hippocampal activity from human epileptic patients.. We observed hippocampal theta waves increased with higher speed and higher sensory demands, thereby maintaining constant information per oscillatory cycle.

These results highlight the role of hippocampal theta oscillations in discretizing ongoing experience relative to the available information and explain how episodic memory integrates a fixed number of items per oscillatory theta cycle irrespective of richness of the external world.

## Introduction

George Miller recognized that the capacity of memory does not have a defined bound but rather is organized around chunks which can be variably defined dependent on experience^1^. This hypothesis emphasizes that the brain overcomes information bottlenecks through recoding and raises the fundamental question of how the brain forms episodic memories from the continuous stream of information that the sensory systems relay. A related issue is what the dynamics and mechanism could be of the temporal segmentation and recoding that chunking assumes. It is known that the medial temporal lobe (MTL) plays a critical role in this process of memory chunking. For instance, subjects with MTL lesions, including the hippocampus, show deficits in the formation and retrieval of episodic memories^2,3^, yet they are still capable of recognizing stimuli that require no context, such as a face^4^ or objects^5^. Electrophysiological recordings of hippocampal neurons show that single units can encode unique locations within a room^6^, with pools of neurons producing cognitive maps that can cover the whole environment^7^. This discretized representation of space, in turn, can support prospective processes such as trajectory planning^8^ and mental time-travel^9^. Mnemonic processing is not restricted to spatial information alone as hippocampal neurons can also encode time intervals^10^, objects^11^, other agents^12,13,^ and the identity of different rooms^14^, illustrating the multi-modal and abstract nature of episodic memory^15,16^. As an example, rodents can ‘navigate’ through a soundscape displaying hippocampal responses that overlap with their place and grid cells^17^. Moreover, the phenomenon of rate remapping observed in experiments where the spatial layout of the environment is morphed can be explained in terms of the multiplexing of action and perception in the activity of single neurons in the hippocampus^18,19^. Hence, the hippocampus is a central structure underlying the temporal segmentation and binding of mnemonic processing and the formation of chunks.

Hippocampal dynamics display distinct oscillatory rhythms with the most prominent power observed in the theta (2-8 Hz) and gamma-frequency (30-100 Hz)^20,21^. It is believed that this so-called, theta-gamma code represents discrete items by ensembles of neurons firing in the gamma range, which in turn are sequentially ordered in the theta cycle^22^. Indeed, disruption of theta or gamma oscillations produces distinct memory deficits^23–25^. In addition, it has recently been shown that episodic memory is critically dependent on the dynamic coordination between the hippocampus and Lateral Temporal Cortex (LTC) in the gamma range^26^. Here we propose that the density of input information modulates this distinct relation between the theta and gamma oscillations underlying episodic memory. We contend that in this way, the hippocampus resolves the trade-off between the segmentation and compression of continuous input streams of various complexity and the finite capacity of episodic memory.

Hippocampal gamma-frequency oscillations are generally associated with processes in local neuronal circuits^27,28^, whereas theta-frequency oscillations are associated with more global processes^29^. For instance, theta-frequency oscillations have been shown to systematically vary with self-motion in both non-human mammals^30,31^ and humans^32–34^. Furthermore, it has been observed that the oscillatory dynamics of human hippocampal theta is modulated by the speed of movement during ambulatory behavior^35^. We reinterpret this speed-dependent modulation as a signature of a more fundamental process: the information density-dependent temporal segmentation and compression of continuous inputs streams into episodic memory chunks. Indeed, locomotion speed directly affects both velocity cues, including optic flow, and the amount of encountered sensory cues. We propose that the modulation of theta is actually driven by the aggregated information density of inputs reaching the hippocampus and how they drive hippocampal circuits, irrespective of their cause (self-motion or environmental statistics). This effect, in turn, provides for the information density-dependent normalization of the content of memory chunks. To test this hypothesis, we quantified the contribution of movement speed and the number of visual cues to hippocampal oscillations in the intracranial EEG data acquired from pre-surgical drug-resistant epilepsy patients (Fig.1A-C). We analyzed the relationship between gamma and theta oscillations to show that the hippocampal cross-frequency code is subject to information density-dependent normalization through theta oscillations. As a signature of the information content of episodic memory, we considered P-episodes, which are sustained high-frequency events previously linked to mnemonic encoding^36^. Their distribution along the phase of hippocampal theta can thus be considered as a compression metric. To elucidate the neural mechanisms underlying the memory normalization, we showed with a hippocampal model that theta-gamma nested oscillations display the same theta frequency adaptation and gamma episodes per theta cycle distribution when exposed to increasing levels of information density. This suggests that the memory normalization we report is an intrinsic property of hippocampal circuits.

**Figure 1.**
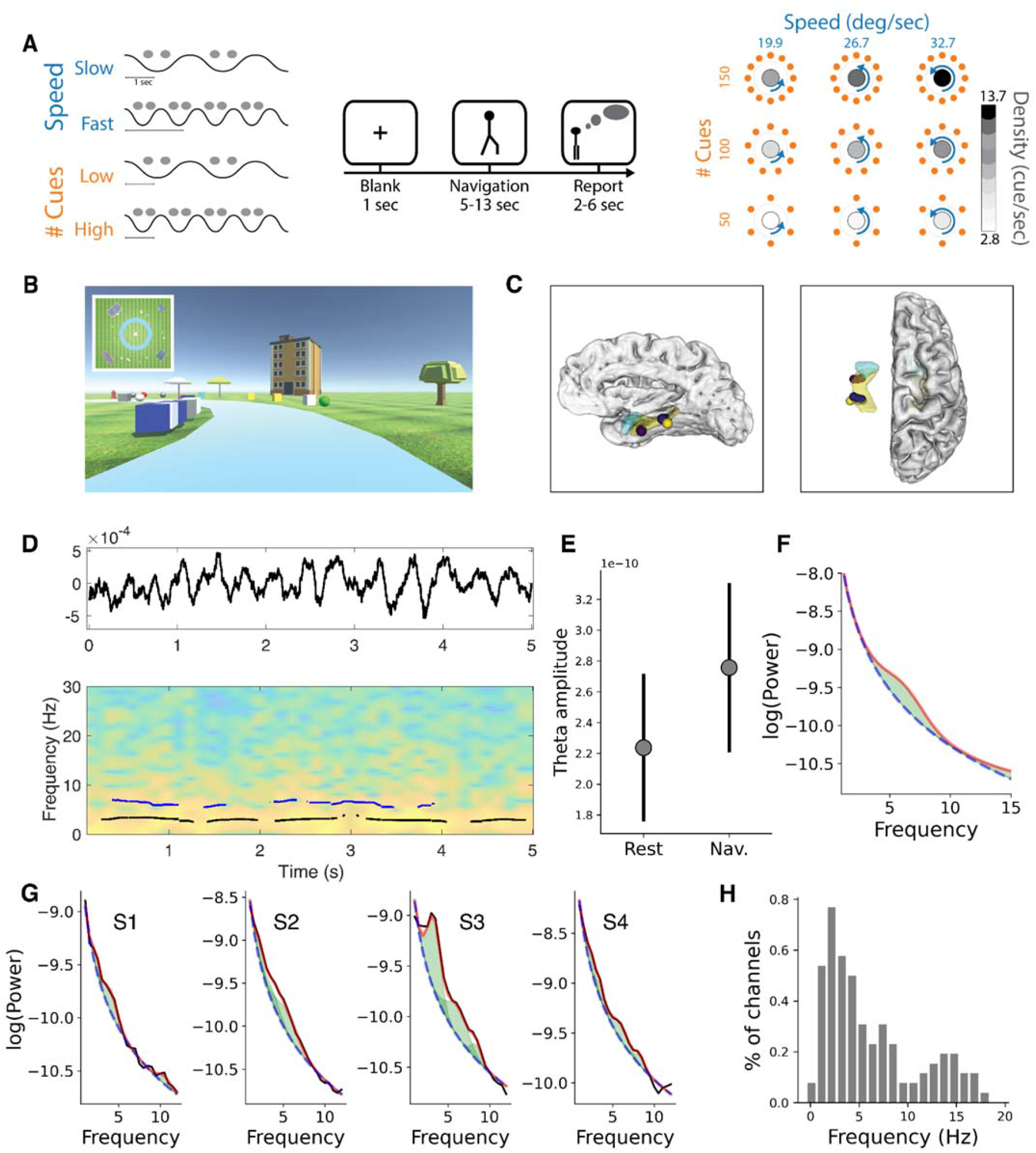
Experimental setup and the effect of information density on theta power. **A-left:** Illustration of the concept of normalization. The speed of locomotion and amount of sensory spatial information modulate hippocampal theta oscillations to accommodate more information encoded in the gamma range. **A-center:** Experimental protocol: each trial starts with a 1 second presentation of a blank screen alerting the subject of the beginning of a new trial. It is followed by a virtual navigation period in first-person perspective lasting from 5 to 13 seconds, and finally by a reporting period lasting between 2 to 6 seconds. **A-right:** Experimental conditions. At each trial, the speed of (virtual) locomotion, as well as the number of environmental cues, were systematically modulated. A third variable was derived from the combination of locomotion speed and the number of cues resulting in the compound measure of number of cues per second or information density. **B:** The virtual environment used for the navigational task (see Methods). The top left inset shows a bird’s eye view of the circular virtual track (shown here only for illustration purposes). **C:** Electrodes location in MNI space coordinates showing their overlap with hippocampal regions (yellow mesh). For reference purposes, amygdaloid volumes are shown in light-blue. **D:** Example of a raw hippocampal LFP (top) and the corresponding spectrogram. Overlaid lines represent the instantaneous frequencies of the detected oscillatory bands when crossing the 1/f local fit. **E**: Increased theta strength during navigation epochs compared to resting (inter-trial) periods (paired T-test, statistic=-2.93, p=0.06, error bars indicate S.E.M.). **F**: Averaged Power Spectral Density (PSD) during navigation. Shaded regions indicate density above 1/f fit. **G**: Same as F for individual subjects. **H**: Distribution of oscillatory peaks detected per recorded channel shows a strong presence of lower frequencies.

## Results

We analyzed the local field potentials (LFPs) from hippocampal iEEG electrodes of four subjects (one female, mean age 34±7) with pharmaco-resistant epilepsy, performing a passive virtual reality spatial navigation task where they moved along a circular track lined with global and local visual cues (Fig.1B). Participants viewed the track from a first-person perspective similar to previous experiments^37^. The global and local cues were placed at uniform distances to the navigational path, and their location and appearance were randomized for each trial. In this way, we were able to manipulate the speed of locomotion, the number of visual cues and their combination: information density (cues per second). The within-subject design had three levels per independent variable, leading to a total of 9 conditions, which allowed us to dissociate the effect of movement speed from that of the number of visual cues (Fig.1A).

To dissociate the effects of locomotion speed and sensory input sampling on the modulation of hippocampal theta waves, we analysed hippocampal local field potentials obtained from the iEEG recordings. Visual inspection of hippocampal raw traces and multiple oscillations detection analises^38^ revealed the expected presence of theta oscillations in the recorded dataset (Fig. 1-D). Consistent with the literature^39^, we observed higher power amplitudes in the power of the theta frequency band (2-8 Hz) during the virtual navigation than at rest, confirming previous observations in animal models and suggesting engagement of the subjects in the task (Fig. 1E). To assess the genuine presence of theta waves during virtual navigation, we parameterized the power spectra of each recorded channel and detected periodic components rising above the 1/f aperiodic fit^40^, and found the majority of channels yielded components in the theta band (Fig. 1F-H and Fig. 1 Supplement 3).

We then analyzed how locomotion speed and the number of visual cues independently affect the dominant frequency of hippocampal theta oscillations. Whereas locomotion speed did not significantly affect dominant theta frequency (Friedman test: χ^2^=3.5, p=0.17, df=2; low vs high speed paired t-test: t=-1.39, p=0.25, df=7, Fig. 2A left and Fig. 3 Supplement 3), the number of cues (Friedman test: χ^2^=6.5, p=0.038, df=2, low vs high number of cues paired t-test: t=-6.29, p=0.008, df=7, Fig. 2A right and Fig. 2 Supplement 3), and information density (cue/sec, Friedman test: χ^2^= 12.2, p=0.015, df=4, low-high cue/sec paired t-test: t=-6.71, p=0.006, df=7, Fig. 2B and Fig. 2 Supplement 3, Fig. 2 Supplement 4), all triggered specific increases in the frequency of theta oscillations.

**Figure 2.**
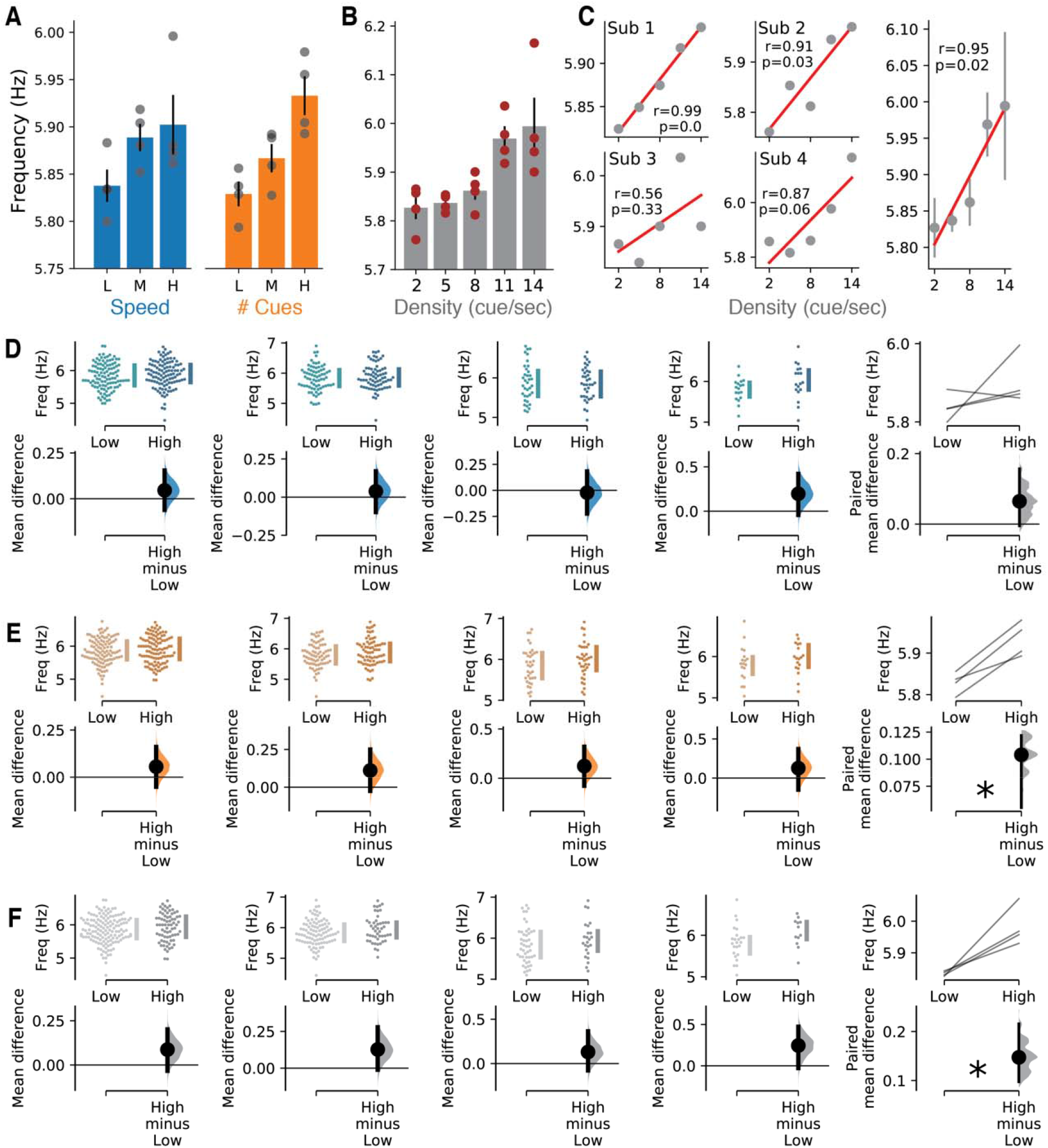
Theta dominant frequency is modulated by information density. **A:** Average dominant theta frequency peaks tend to increase with higher locomotion speed (Friedman Chi-square: statistic=3.5, p=0.17; Low-vs-High speed paired T-test: statistic=-1.3, p=0.2), and larger number of environmental cues (Friedman Chi-square: statistic=6.5, p=0.038; Low-vs-High number of items paired T-test: statistic=-6.2, p=0.008). **B:** Exposure to higher information density leads to increases in theta frequencies (Friedman Chi-square: statistic=12.2, p=0.01; 2-vs-11 cue/sec paired T-test: statistic=-6.7, p=0.006). **C:** Individual subjects and population averaged correlations between theta frequency and cue/sec condition (labeled Pearson-R for each individual). **D, E, and F:** Individual and population level bootstrap density analysis confirms effect of number of environmental cues and information density conditions in modulating theta frequency. **D** Speed (mean difference=0.06; bias corrected confidence interval = -0.004,0.15; Student t-test: p=0.2, t=-1.39). **E** Number of environmental cues (mean difference=0.1; bias corrected confidence interval=0.05,0.12; Student t-test: p=0.008, t=-6.29). **F** Information density (mean difference=0.14; bias corrected confidence interval = 0.09,0.21; Student t-test: p=0.02, t=-4.33).

The positive relationship between dominant hippocampal theta frequency and information density was further observed by correlation analysis within and across subjects (Pearson-r test: r=0.95, p=0.02, df=3) (Fig. 2C). In addition, we estimated the effects of speed, number of cues, and information density on hippocampal theta by comparing their low and high levels for each subject and the aggregated population using bootstrap-coupled estimation analysis, which confirmed the effect of information density in modulating theta rhythms (Fig. 2E-F).

Because increases in both locomotion speed and the number of cues increases lead to higher information density, the observed increase in theta frequency may be a consequence of eye saccades triggering oscillatory resets as observed in non-human primates^41^. To control for this alternative interpretation, we quantified individual cycle symmetry andperiod consistency for each subject to identify abrupt changes in theta phase^42^. We found no significant difference across the experimental conditions thus supporting our interpretation that the hippocampus normalizes information content of memory (Fig. 1 Supplement 1, Fig. 1 Supplement 2, Fig. 2 Supplement 1).

The theta-gamma coding scheme alludes to a systematic phase-amplitude relationship between the two frequency bands, where gamma displays higher power at a specific phase interval^22^. To assess how the hippocampal theta-gamma code is affected by information density, we calculated the distribution of gamma amplitudes (40-80 Hz) along with the theta phase (Fig. 3A-C) by calculating the theta-gamma distribution against a uniform distribution (modulation index)^43,441,42^. Irrespective of the information density load, we observed higher gamma amplitude in the third theta cycle quadrant (π) (Q3(π) comparison; Paired t-test, Q1: t=10.80, p=0.0004, df=9, Q2: t=3.32, p=0.02, Q4: t=3.07, p=0.03, df=9, Fig. 3A).

**Figure 3.**
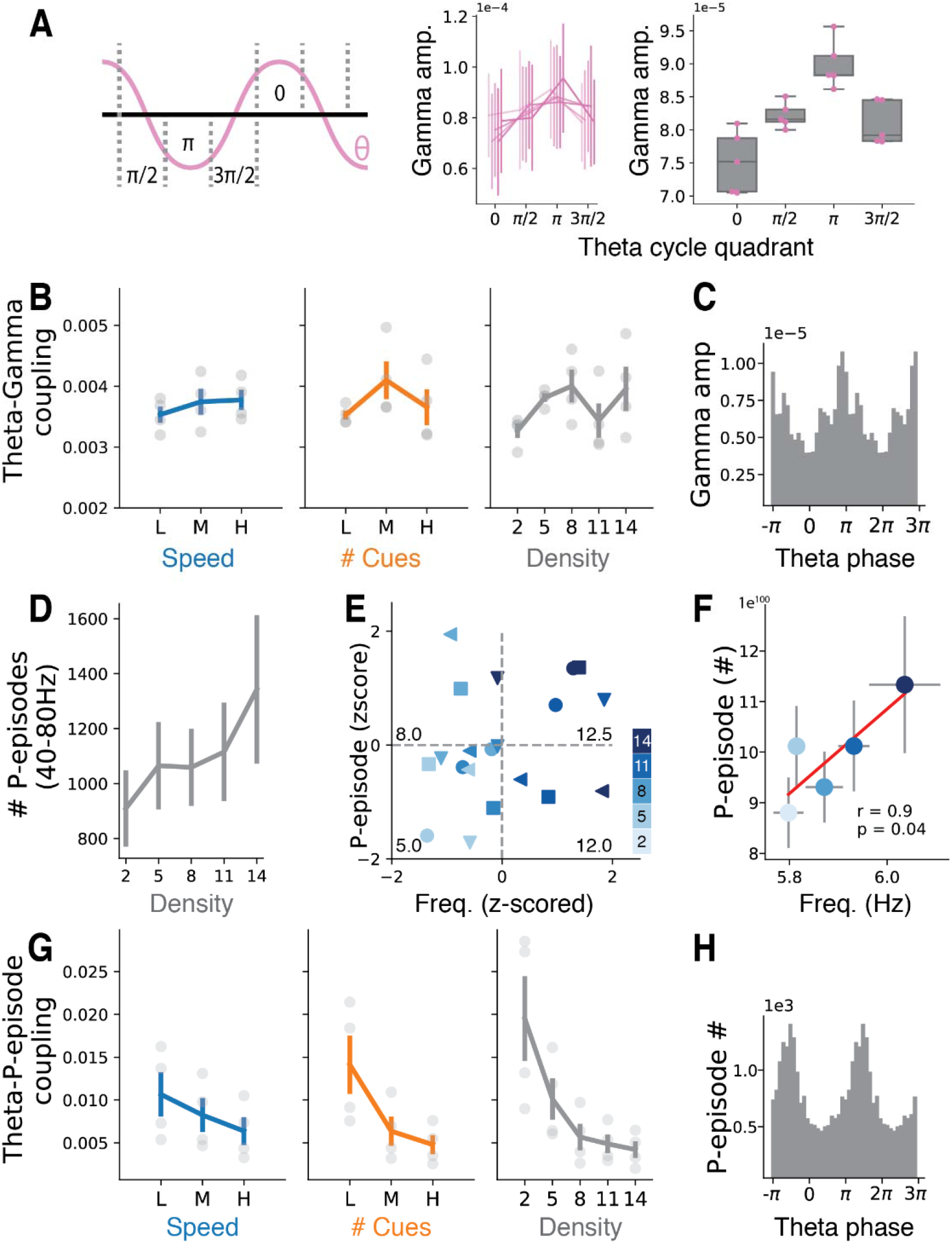
Gamma responses versus information density **A:** Locking of gamma amplitude to theta cycle quadrant. Left: Cartoon representation of one theta cycle and its corresponding quadrant-split. Center: Averaged hippocampal gamma amplitude of all subjects along the four theta quadrants for all information densities conditions (bright-to-dark pink gradient representing from low-to-high information density level). Right: Same as center, with grouped information density levels along the theta-quadrants. Note the persistently stronger gamma amplitudes at the third quadrant (Q3(π)) (Q3(π) comparison; Paired t-test, Q1: t=10.80, p=0.0004, Q2: t=3.32, p=0.02, Q4: t=3.07, p=0.03). **B:** Theta-gamma coupling (modulation index) is not modulated by speed (Friedman Chi-square: statistic=3.5, p=0.1), number of environmental cues (Friedman Chi-square: statistic=2.0, p=0.3), or information density (cue/sec) (Friedman Chi-square: statistic=-1.8, p=0.15). **C:** Distribution of gamma (40-80Hz) amplitudes along with theta (2-8Hz) phase for a representative subject (ID 3). **D:** The number of P-episodes increases along with information density (Friedman Chi-square: statistic=16.5, p=0.035; Low-vs-High density paired T-test: statistic=-2.21, p=0.06). **E:** Distribution of the normalized (z-scored) P-episodes count and theta frequency for trials of distinct information densities (2 to 14 items/sec, color bar). Each marker type is the average of one subject per information density (cue/sec) condition (Pearson-R, r=0.3, p=0.2). Note the increases in the normalized (z-scored) number of P-episodes and theta frequency accompanies the cue/sec level from which each data point was derived from (mean 5cue/sec lower-left, mean 12.5cue/sec upper-right). **F:** Averaged P-episode and dominant frequency across subjects along distinct information density conditions (see colorbar in **E**). Horizontal and vertical error bars represent S.E.M for frequency and P-episode, respectively (Spearman’s rank correlation, r=0.9, p=0.04). **G:** Strong modulation of Theta-P-episode coupling for speed (Friedman Chi-square: statistic=8.0, p=0.01; Low-vs-High speed paired T-test: statistic=4.12, p=0.02), density (Friedman Chi-square: statistic=8.0, p=0.018; Low-vs-High density paired T-test: statistic=4.0, p=0.027), and cue/sec (Friedman Chi-square: statistic=3.8, p=0.003; 2-vs-14 cue/sec paired T-test: statistic=3.81, p=0.03). **H:** Example distribution of P-episode count along with theta (2-8Hz) phase for subject 3.

We did not find significant changes in the modulation index for either condition in speed (Friedman, stat=3.5, p=0.17, df=2), number of cues (Friedman, stat=2.0, p=0.36, df=2), or information density (Friedman, stat=5.6, p=0.23, df=4), suggesting that the positive correlation between information density and dominant theta frequency promotes a robust theta-gamma coding irrespective of information density (Fig 3B and C, and Fig. 3 Supplement 2).

To quantify the impact of the information density-dependent modulation of theta on sensory encoding itself, we determined the occurrence of, P-episodes, that is the transient oscillatory events of high power in the gamma band reflecting the processing of sensory information. We observed a systematic increase in the number of P-episodes with the increase of information density (Friedman Chi-square: statistic=16.5, p=0.035, df=4; Low-vs-High information density paired T-test: statistic=-2.21, p=0.06, df=7, Fig. 3D), suggesting that the latter leads to an increase of sustained gamma activity.

Thus far, our results show information density-dependent modulation of theta oscillations and P-episodes with a constant phase relationship between theta and gamma frequencies. This favors the idea that hippocampal theta oscillations operate as a normalization mechanism encoding a constant amount of information per cycle. This interpretation predicts that an increase in the number of observed P-episodes is accompanied with an increase in dominant theta frequency which is confirmed by our analysis (Fig. 3E and F, r=0.9, p=0.04, df=3, Spearman test). However, contrary to what we observed for theta-gamma coupling, P-episodes decreased in their preferred phase relative to the modulated theta cycle for speed (Friedman Chi-square: statistic=8.0, p=0.01, df=2; Low-vs-High paired T-test: statistic=4.12, p=0.02, df=7), density (Friedman Chi-square: statistic=8.0, p=0.01, df=2; Low-vs-High paired T-test: statistic=-2.21, p=0.06, df=7), and cue/sec (Friedman Chi-square: statistic=15.4, p=0.003, df=4; Low-vs-High paired T-test: statistic=3.81, p=0.03, df=7) conditions (Fig. 3G and H).

To further elucidate the putative mechanisms underlying information normalization in the hippocampus, we built a spiking neural model that combines recurrent fast and slow inhibitory feedback onto excitatory neurons through different types of interneurons^45,46^ (Fig. 4A, top left). We exposed the model to a stimulation protocol resembling the traversal of a linear track driving the sequential occurrence of overlapping asymmetrical place fields^47^ (Fig. 4A, top right). We defined information density as the total number of active place cells. Under these conditions, the model displays nested oscillations reflecting a theta-gamma phase code^48^ (Fig. 4B, E), as shown in the frequency analysis performed to the LFP-like signals and the spike times extracted from the population of pyramidal cells (Fig. 4B). We observe high power in the theta (4-10 Hz) and gamma (30-50 Hz) range (Fig. 4D) across conditions. Additionally, gamma amplitude (30-80 Hz) is significantly modulated by the phase of theta in all conditions (Fig. 4C), ensuring stable episodic representations in theta cycles for all three levels of information density. Moreover, a significant negative correlation is observed between the position along the place field, associated place cell activity and the theta phase (Fig. 4E) indicative of phase precession^49^. Consistent with our physiological results the model displays a positive correlation between information density and theta frequency (Fig. 4F, left) while the number of gamma episodes per theta cycle remains constant (Fig. 4F, right). Hence, these results from our model suggest that the normalization we observe could be attributed to intrinsic hippocampal dynamics.

**Figure 4.**
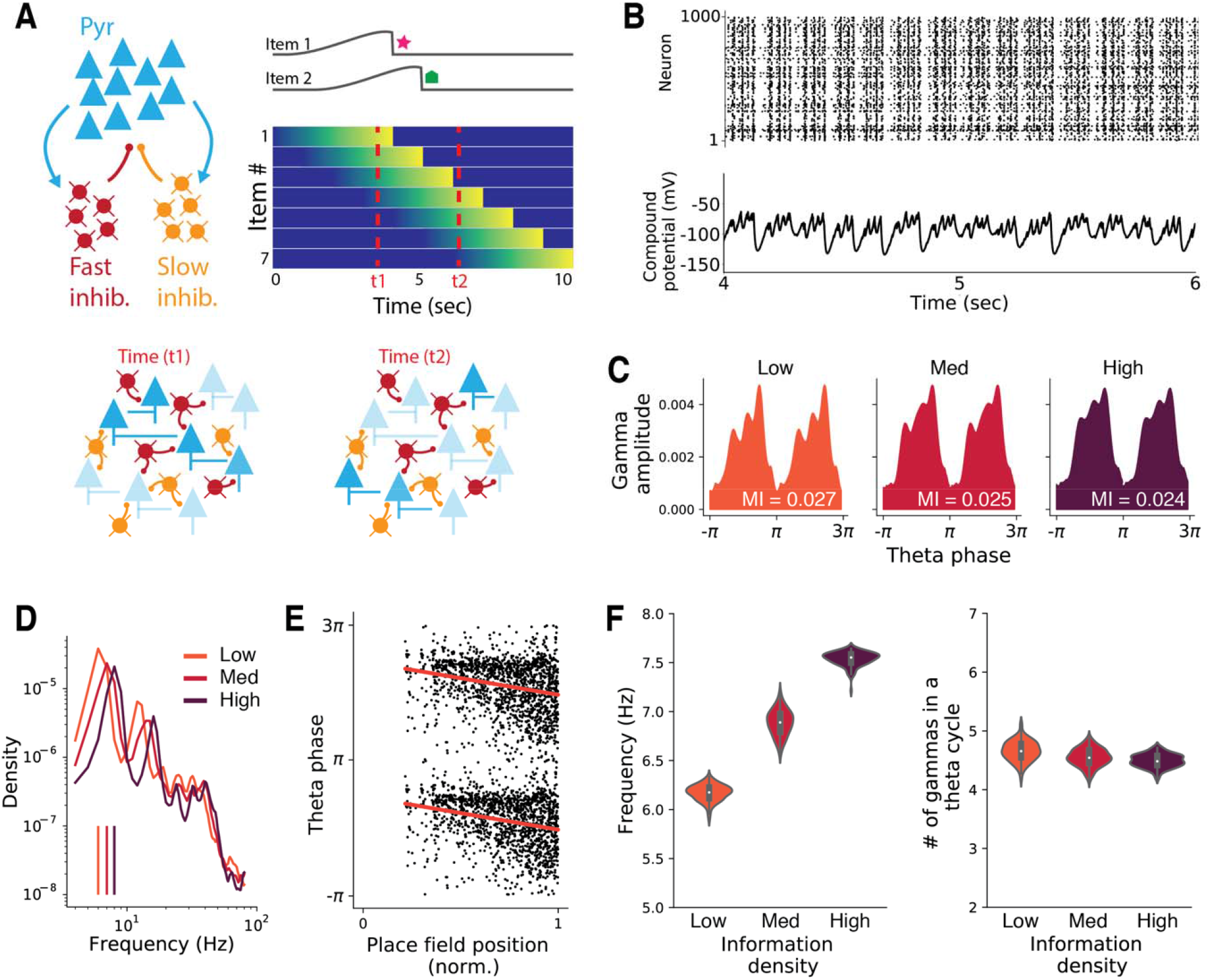
Hippocampal computational model demonstrating the mechanisms underlying information normalization. **A:** Schematic illustration of the model. Pyramidal cells encoding for items receive fast and slow global inhibitory feedback (see Methods). The stimulation protocol is based on asymmetric place fields representing items encountered along a linear track. Each item stimulates a distinct population of pyramidal cells. **B:** Peristimulus time histogram of pyramidal cells with the corresponding LFP-like signal recorded while traversing the linear track. **C:** Theta-gamma phase-amplitude coupling across conditions (p<0.0001 for all MI). MI corresponds to the modulation index, defined as the Kullback-Leibler divergence score against a uniform distribution. **D:** Averaged power spectral density (PSD) across conditions. Theta peaks are indicated with vertical lines. Axes are plotted in a logarithmic scale. **E:** Spikes within the place field and the relative theta phase. Red line represents the linear fit (Pearson r=-0.28, p<0.001). **F:** The relationship of information density with theta frequency (PSD center of mass over 4-8 Hz) and gamma episodes per theta cycle. In all analyses, theta frequency ranges from 4 to 8 Hz, and gamma from 30 to 80 Hz.

## Discussion

We addressed the question of how the continuous stream of information the brain is exposed to during episodic memory is discretized into chunks. We looked at the organization of the oscillatory dynamics of the LFPs in the hippocampus using iEEG recordings obtained from depth electrodes implanted in drug-resistant epileptic patients that were navigating a virtual reality circular maze. By systematically varying the speed of locomotion and the number of visual cues in the environment, we showed that the dominant peak of theta oscillations correlates with overall information density rather than speed alone as reported in the animal literature^50,51^. We also demonstrated that, while the number of P-episodes correlates directly with information density, gamma oscillations remain at a constant phase relationship with theta oscillations. All of these suggest that the theta frequency, gamma amplitude, theta-gamma phase, and P-episodes are dynamical signatures of information content normalization in the memory system.

Our results support the idea that hippocampal theta oscillations realize a discretization of continuous information streams. These oscillations are in turn represented in the gamma range. As activity in the gamma range increases, theta frequency accelerates to reduce the number of gamma cycles that can be encoded per theta cycle, showing that this mnemonic discretization normalizes gamma content relative to overall information density.

Should be noted, this study was conducted in a virtual reality setup and subjects did not actively navigate within a physical environment. The contribution of self-motion signals in modulating hippocampal oscillations is likely weaker than when physical locomotion takes place in real-world environments. Moreover, the presented experimental protocol lacked to assess behavioral measures of spatial situating and environmental statistics, a metric that could further elucidate our understanding of the role of theta oscillations in associative learning and episodic memory formation. Similarly, even though we have assessed the distribution of abrupt changes in the theta phase during navigation, we cannot entirely rule out the potential contribution of eye movements in modulating the relationship between active sampling, hippocampal oscillations and spatial learning. We used a model of the hippocampus to understand the putative mechanisms behind memory normalization which comprised excitatory populations with both fast and slow inhibitory feedback^45,46,52–54^. The specific question our model answered is whether these well-known intrinsic inhibitory feedback loops suffice to account for the normalization effect we observed. Our model unwraps its rate-based inputs in time (interpreted as overlapping place fields), ordering them as gamma episodes in the theta phase consistent with the notion of theta-gamma coding^48^. Furthermore, our model shows that the combined fast and slow negative feedback loops established through inhibitory interneurons accurately account for the online modulation of the theta frequency while maintaining a constant amount of information per cycle carried by the gamma oscillations. This demonstrates that the intrinsic feedback and their associated nested oscillations in the human hippocampus can support the memory normalization that we have discovered.

Miller observed that the capacity of memory does not have a defined bound but rather is organized around chunks which can be variably defined and dependent on experience^1^. The theta-gamma code of the hippocampus shown here provides an illustration of this functionality. By regulating the binding of items in episodic memory relative to information density, the open-ended nature of chunking and its incremental expansion are shown to be dependent on the hippocampus to achieve an efficient re-coding and compression of experiences into memory.

## Figures

**Figure 1-figure supplement 1:**
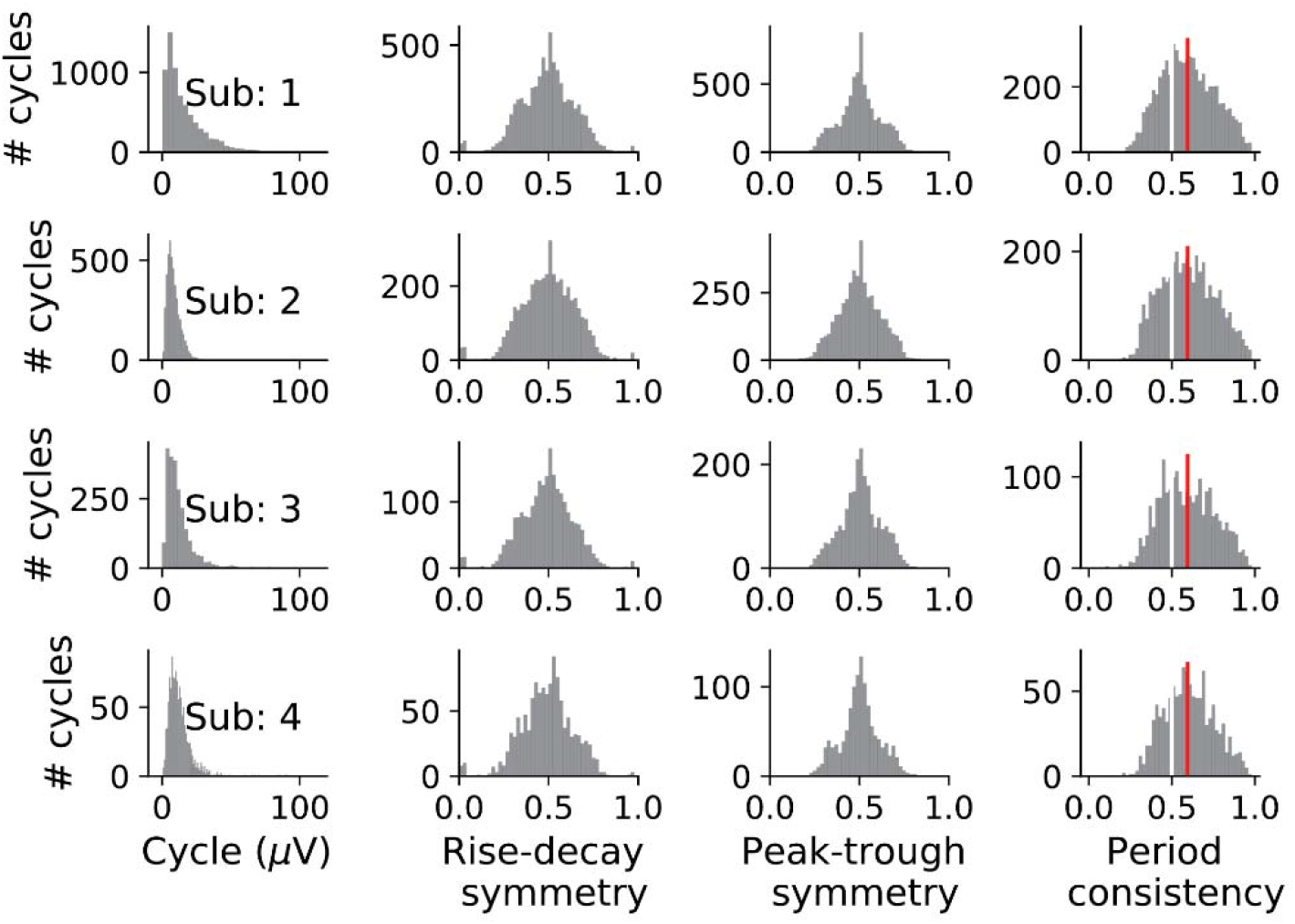
Analytical analysis of subjects’ theta cycles^19^. Note the normal distribution of cycle symmetry and consistency for each participant. Period consistency revealed a skewed theta cycle profile as observed in the rodent hippocampus (see Methods section).

**Figure 1-figure supplement 2:**
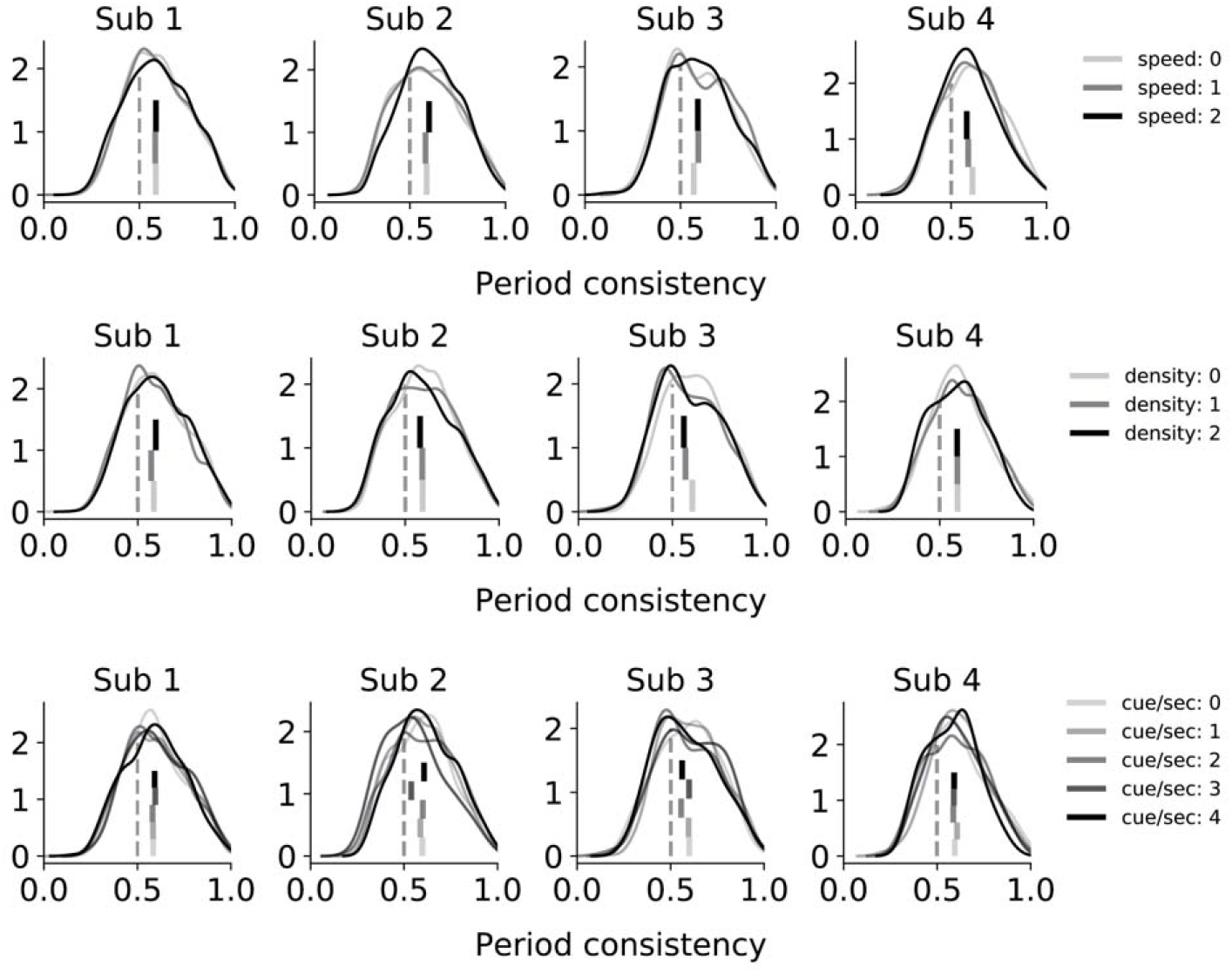
Theta cycles period consistency is not affected by information content^19^. Distribution of period consistency is not affected by the distinct levels of speed, density and cue/sec load, suggesting that individual cycles do not phase-reset.

**Figure 1-figure supplement 3:**
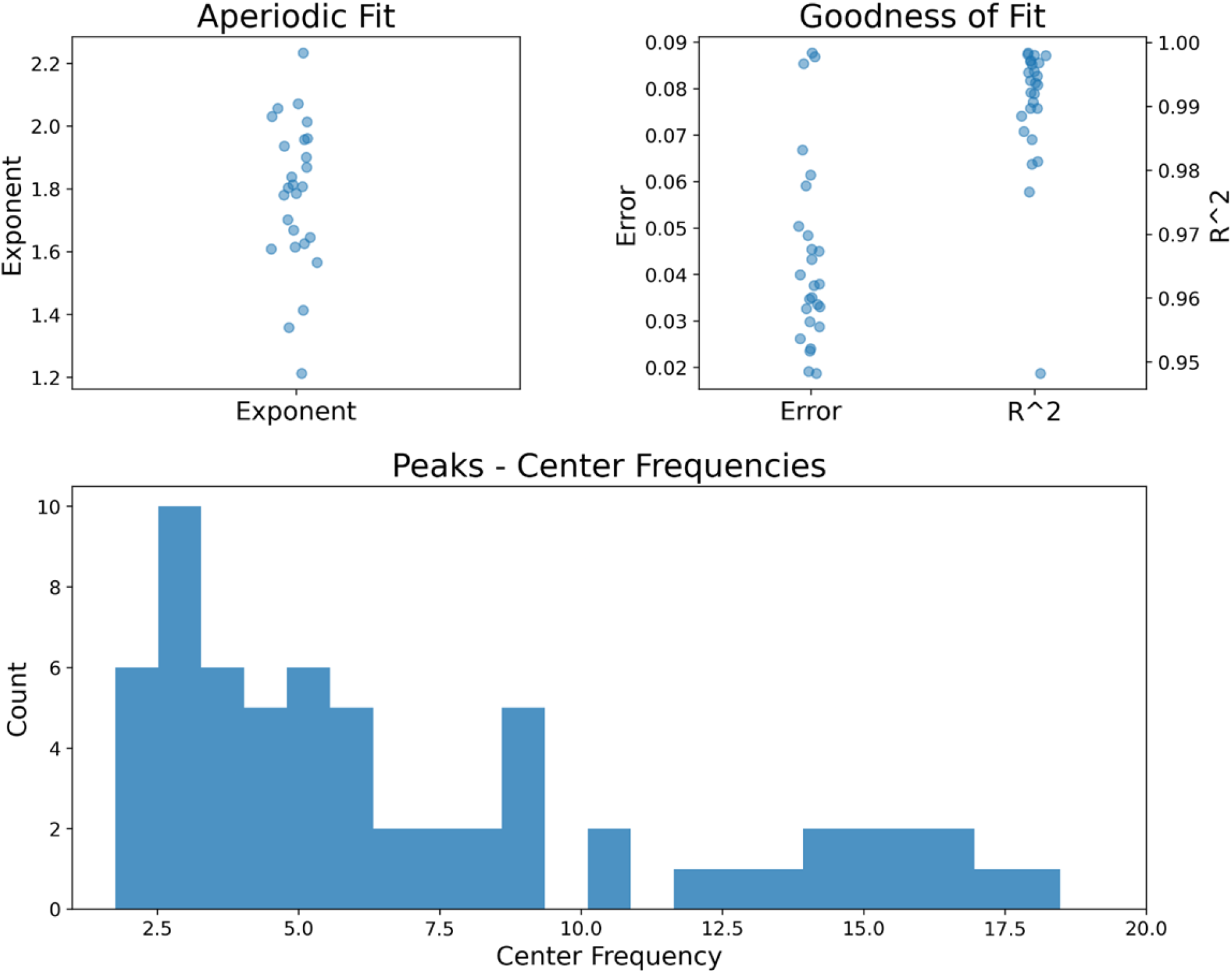
Distribution of oscillatory peaks detected above 1/f local fit of the spectral density for each channel and their goodness of fit.

**Figure 2-figure supplement 1:**
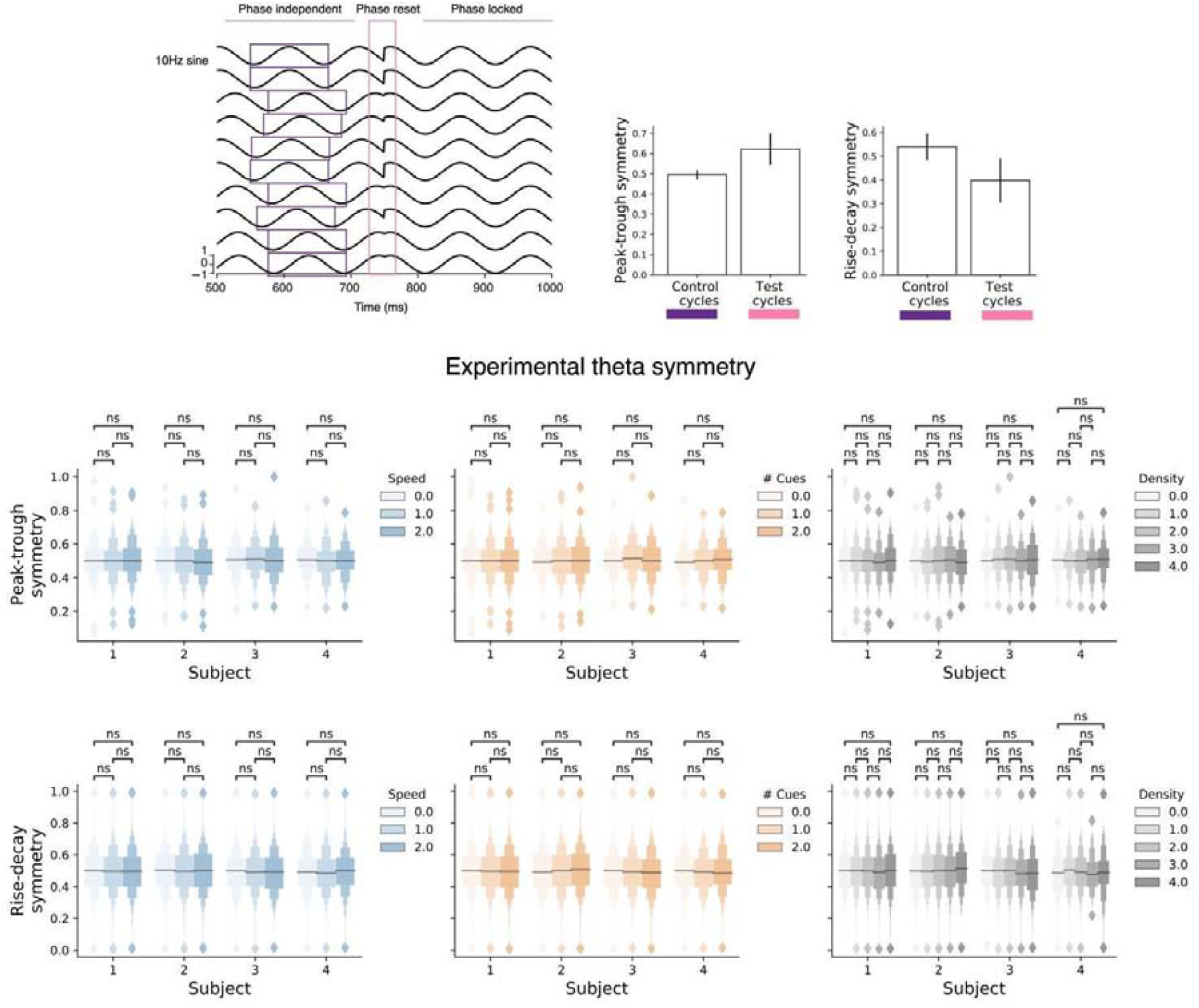
Theta cycle profile for analytical features characterizing an oscillatory signal. **Top** Illustration of the analysis method on artificially generated sine waves to capture changes in oscillatory phase. Sinusoidal waves (10Hz) with distinct phases were concatenated (n=100). Peak-trough and rise-decay symmetry measures were used to assess phase reset in control and test cycles (with 0.5 denoting wave symmetry, see Methods section). **Bottom** To assess the rate of phase reset along our experimental conditions, we used Peak-trough and rise-decay symmetry measures for individual subjects^19^. None of the variables (Speed, # of Cues, or Density) levels (low to high) revealed significantly asymmetric profiles, suggesting that hippocampal theta phase was not greatly affected by eye movements during the experimental task.

**Figure 2-figure supplement 2:**
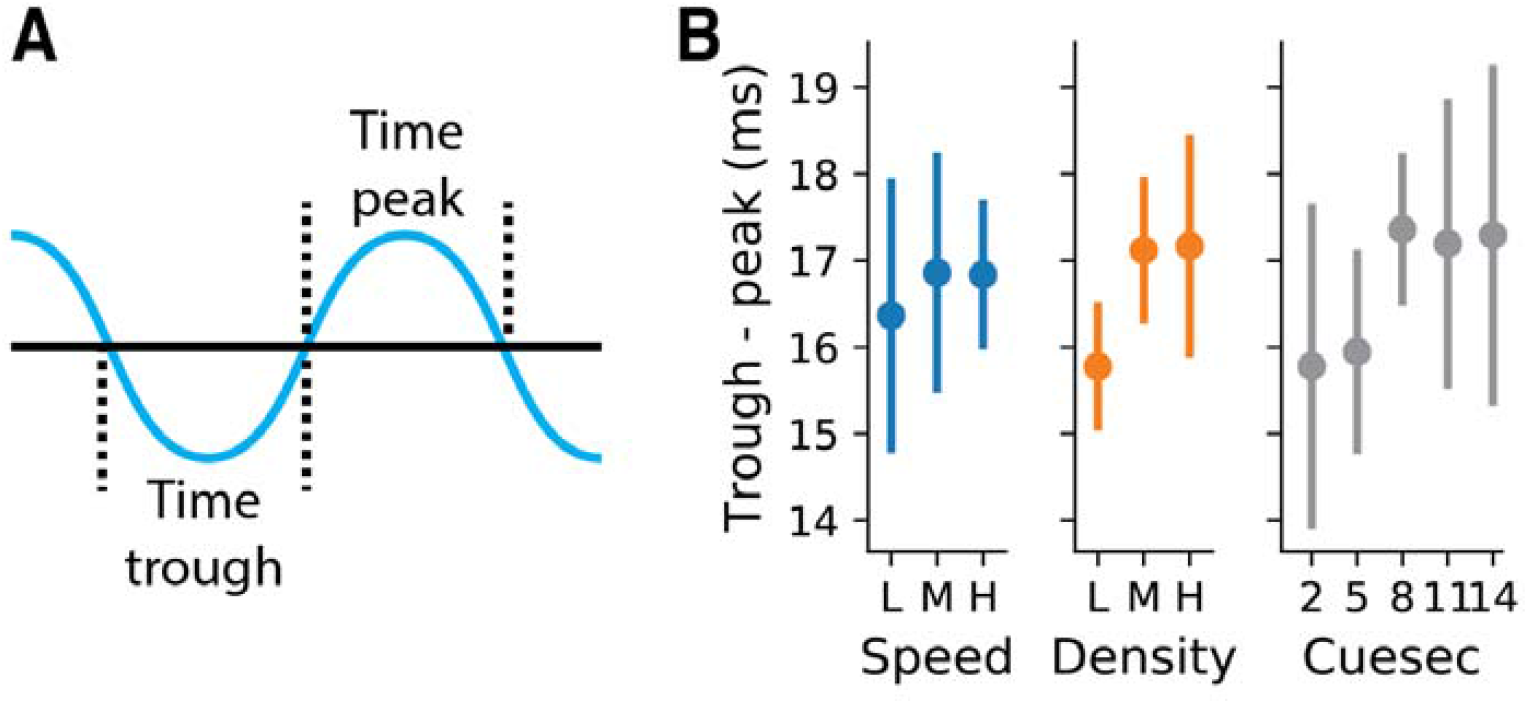
Increases in the relative theta cycle skewness (Time trough – time peak) accompanying the increases in density (Friedman, stat=6.0, p=0.04; T-test low vs high, t=-2.7, p=0.06) and cue/sec (Friedman, stat=5.0, p=0.2; T-test low vs high, t=-3.5, p=0.03), but not for speed (Friedman, stat=0.5, p=0.7; T-test low vs high, t=-0.7, p=0.5) suggesting that environmental sensory information has a higher contribution in modulating hippocampal theta oscillations compared to locomotory-related signals.

**Figure 2-figure supplement 3:**
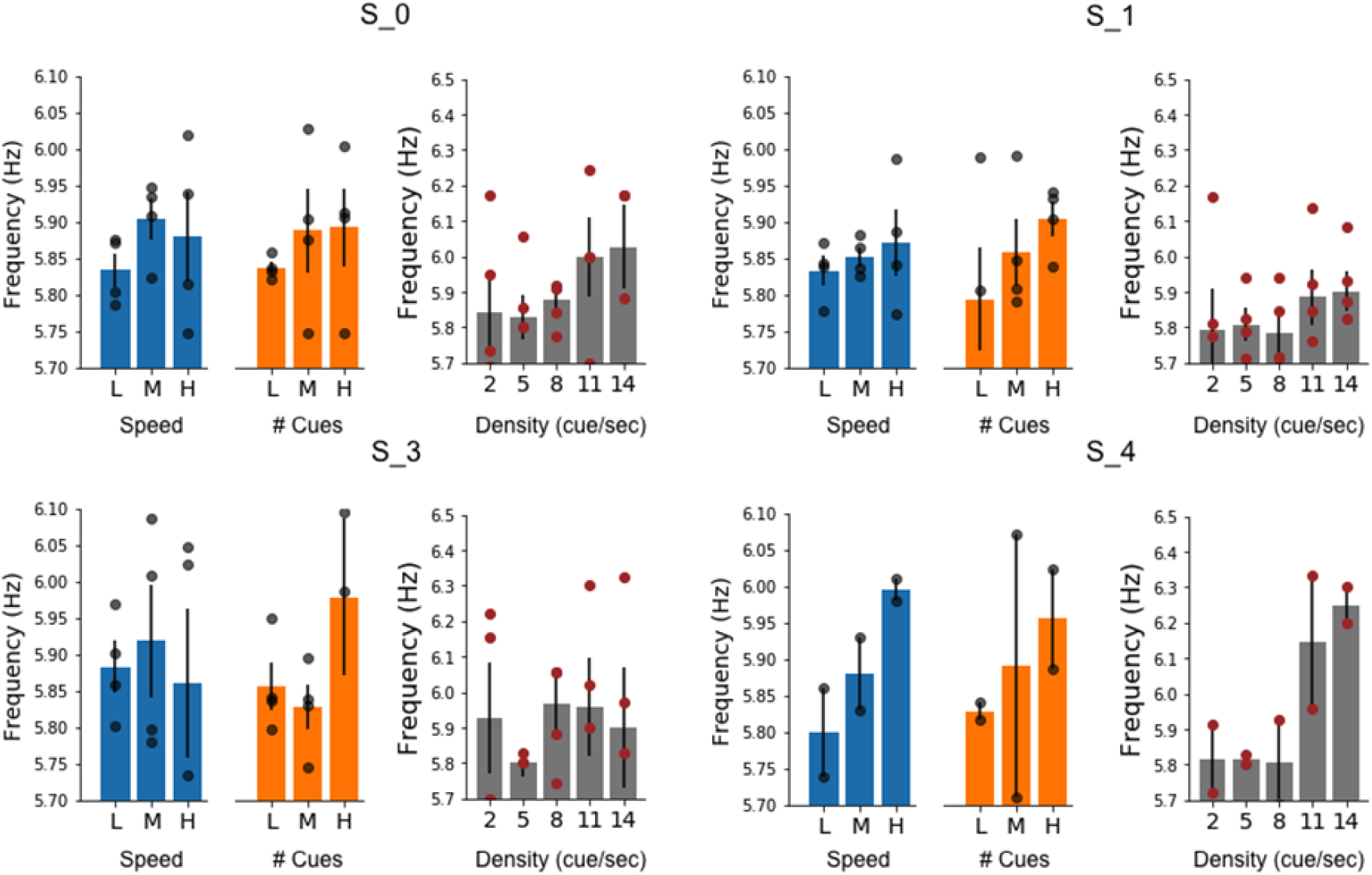
Modulation of the theta dominant frequency by information density as reflected by the individual contact points for each subject.

**Figure 2-figure supplement 4:**
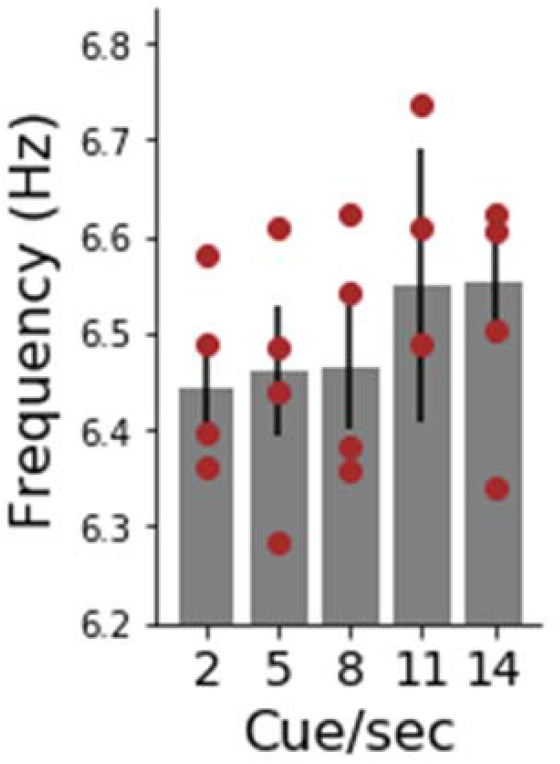
Upward modulation of theta frequency by information density as demonstrated by assessing the instantaneous frequency shifts.

**Figure 3-figure supplement 1:**
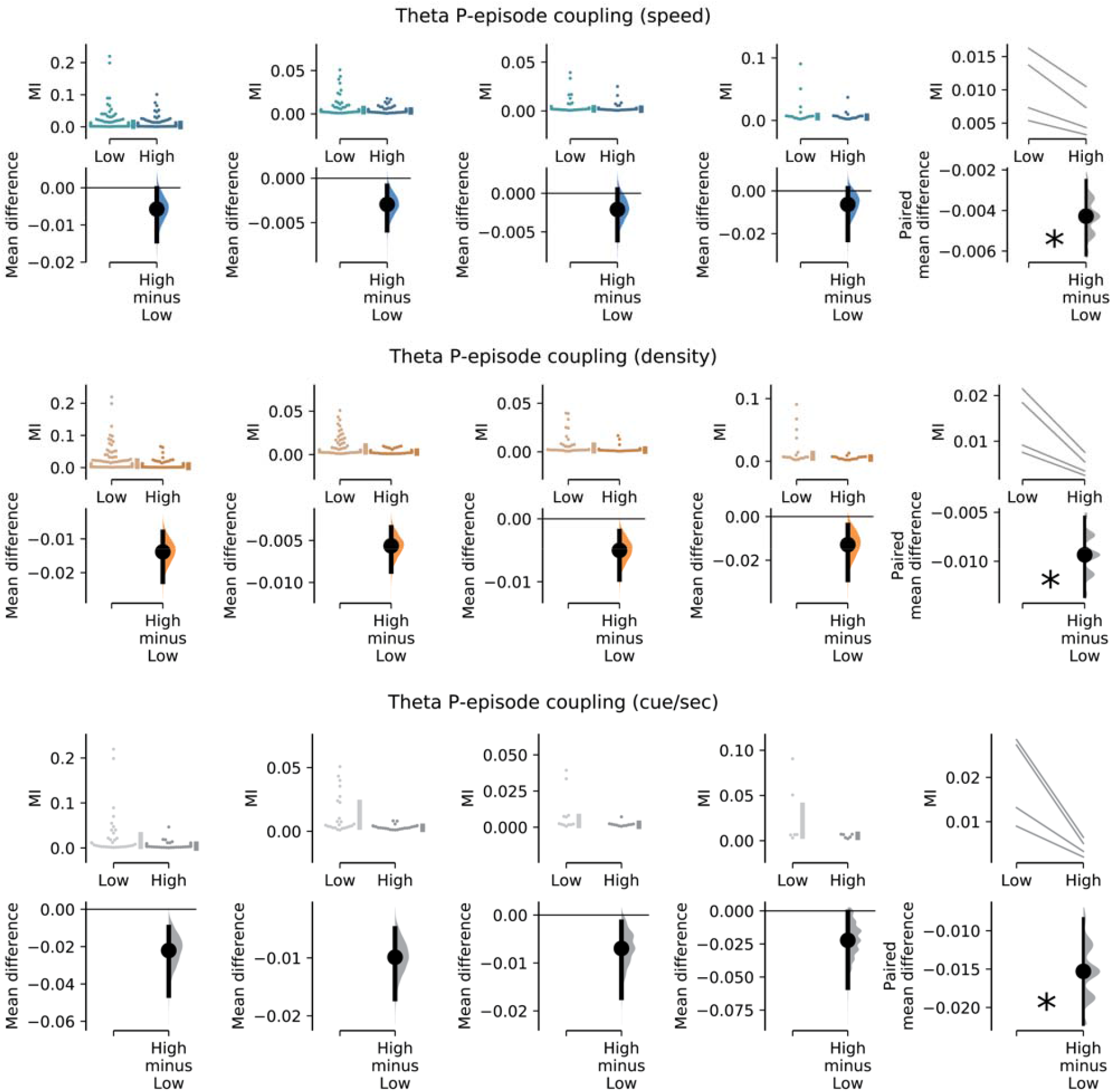
Systematic decreases in Theta P-episode coupling (M.I.) for population and single participant analysis for the distinct levels of speed, density and cue/sec load.

**Figure 3-figure supplement 2:**
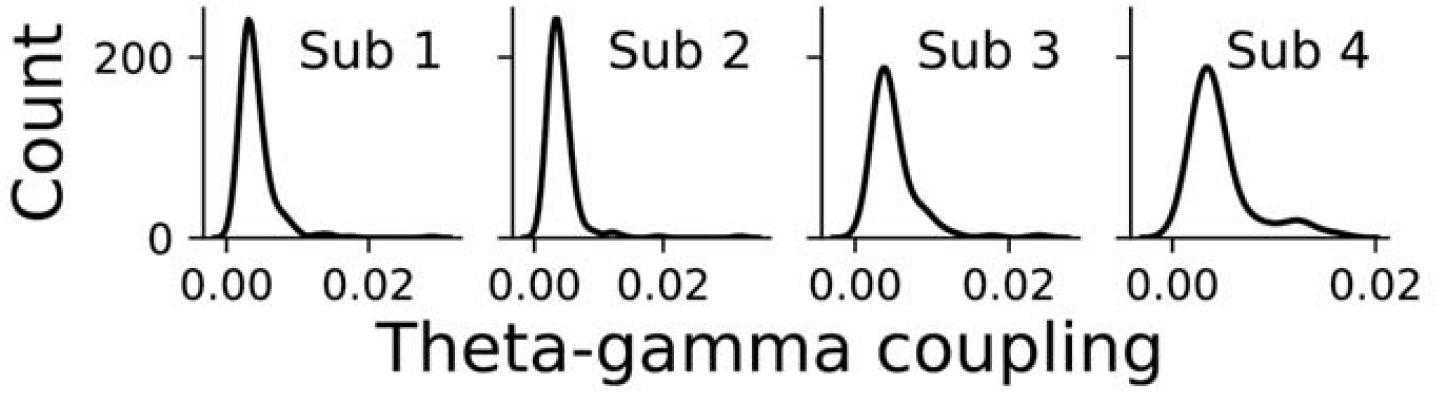
Theta-gamma coupling distribution for individual subjects.

## Methods

### Participants

Four medically refractory epilepsy patients (1 female, mean age 34±7 years old), evaluated for presurgical diagnosis in the Epilepsy Monitoring Unit of the Hospital del Mar (Barcelona, Spain), participated in this study. Patients had been surgically implanted with depth electrodes for diagnostic screening, preceding the surgical treatment. All patients underwent a neuropsychological evaluation and had a normal or corrected-to-normal vision. Patients’ demographic and clinical data are provided in Table 1. The study has been approved by the “Clinical Research Ethical Committee (CEIC), Parc de Salut Mar” (Barcelona, Spain) and, following the Declaration of Helsinki, all participants were informed about the procedure and provided written consent before participating in the experiment.

**Table 1.**
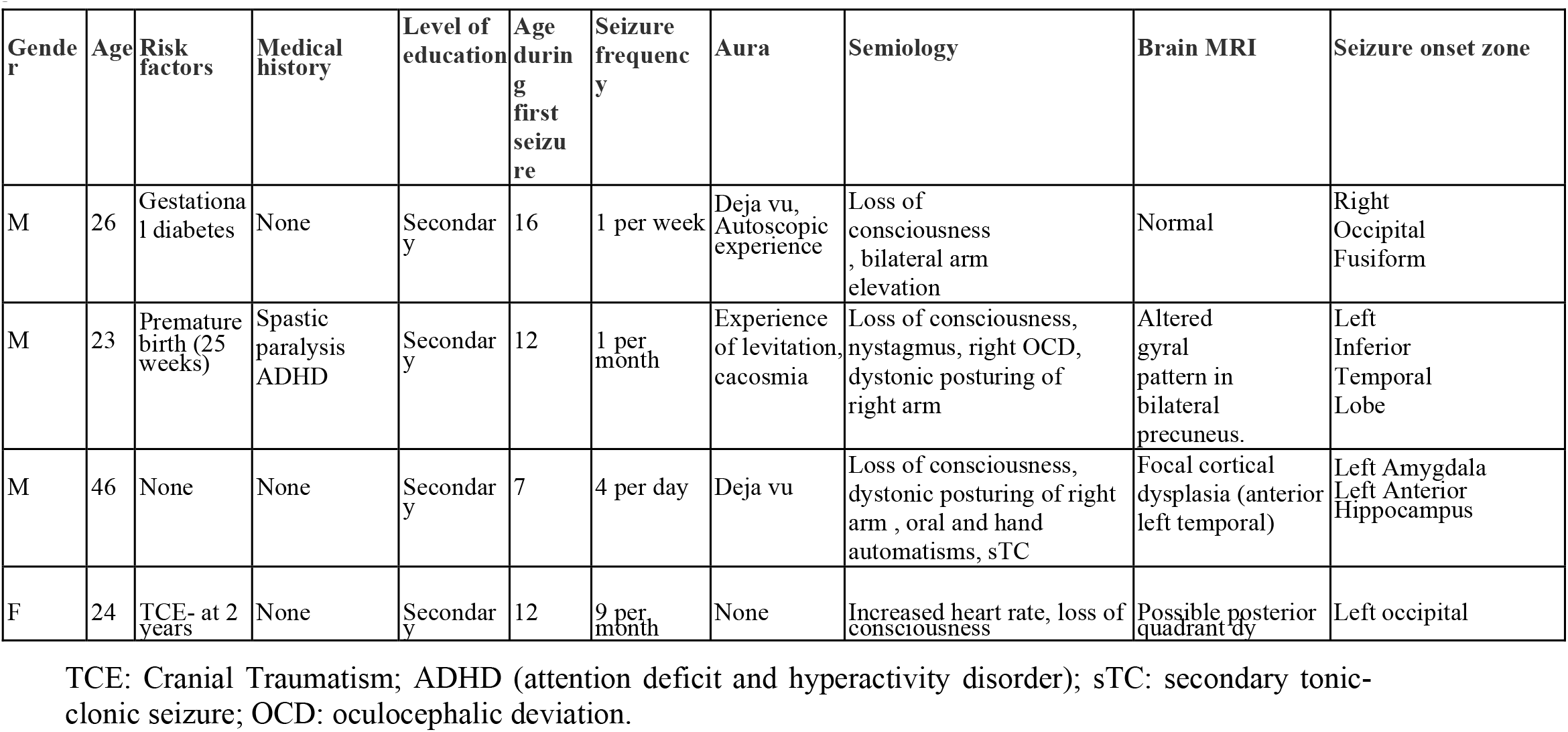
Patients’ demographics and clinical data

### Behavioral task

Participants performed a passive navigation task in a Virtual Reality (VR) environment within a circular track at a constant speed. Four 3D models of buildings placed on the outer part of the track (one per quadrant) served as global environmental landmarks during navigation. Within the track, a set of local cues composed of everyday-life objects, obtained from the “texture city prefabs” library for Unity 3D (https://assetstore.unity.com/packages/3d/environments/urban/1texture-city-112740), were randomly selected and placed at uniform distances along the navigational path (see figure 1B for a snapshot of the virtual environment). Similarly to (1), participants viewed the track from a first-person perspective. The starting point position in each trial was always the same and participants were facing the same direction. Patients participated in either one (N=2), two (N=1), or three experimental sessions (N=1). The second and the third sessions took place on consecutive days. Each session consisted of 27 trials, and each trial consisted of a navigation traversal along the circular track lasting for 5-13 seconds depending on the experimental condition. At each trial, the number of local landmarks placed in the surroundings of the circular track as well as the locomotion speed of the virtual avatar varied accordingly with the experimental condition. The number of visual cues placed on the track was set to 50, 100, or 150 landmarks, whereas the speed of locomotion was defined as 19.9, 26.7, or 32.7 deg/sec, for the low, medium, and high information density conditions respectively. The trial’s order was fully randomized. Each trial started with a 1-sec blank screen with a black fixation cross in the center alerting the subject of the beginning of a new trial. At the end of the navigation, a blank screen appeared for 4 seconds, and participants were probed by asking to report about the distribution of local landmarks verbally. The verbal reporting was used to promote engagement with the task; however, no quantification of performance was carried out.

Patients performed the task on a 15” Apple MacBook laptop while sitting in a comfortable position in their hospital bed at a distance of approximately 60 cm from the screen. The VR application was created using the Unity3D game engine (Unity Technologies, San Francisco, CA, USA).

### Electrodes selection and localization

Stereotactic implantation of depth electrodes was performed by the clinical team at Hospital del Mar (Barcelona, Spain). The location of the electrodes was established only for clinical reasons using an SEEG approach. Targeted regions varied across patients depending on their clinical assessment, but in all participants recruited for this study, they included the hippocampus in the left (n=3) or the right hemisphere (n=1). After co-registering pre- and post-electrode placement using MR scans and CT whole-brain volumes, we could confirm six contacts located in the anterior hippocampus and eight contacts in the posterior hippocampus (see Figure 1C). Locations of the electrodes in native space were finally converted to the standard Montreal Neurological Institute (MNI) space using 3D Slicer (2) and BrainX3 (3), following the method in (4).

### Electrophysiological recordings

Recordings were performed using a standard clinical EEG system (XLTEK, subsidiary of Natus Medical Inc.) with a 500 Hz sampling rate per channel. Unilateral implantation (left hemisphere=3, right hemisphere=1) was performed in all patients using 7 to 10 intra-cerebral electrodes (Dixi Médical, Besançon, France; diameter: 0.8 mm; 5 to 15 contacts, 2 mm long, 1.5 mm apart) that were stereotactically inserted using robotic guidance (ROSA, Medtech Surgical, Inc, Montpellier, France). SEEG signals were re-referenced to the average reference of the same electrode array. Bipolar signals were also derived by differentiating neighboring electrodes pairs from the same electrode array to check for consistency of the results. For all the subsequent analysis, the continuously recorded signal at the selected contact points was initially band-pass filtered in the range 1-200 Hz using a two-way, zero phase lag finite impulse response (FIR) filter to prevent phase distortion. Additionally, to remove the effects of power line noise, a notch filter was applied at 50 Hz and harmonics (fourth-order, 2 Hz bandwidth). Event-triggered TTL pulses were also recorded for synchronization with task events.

### Data analysis

All analyses were performed using custom programming scripts written in Python and using the *mne* library (5), an open-source toolbox for the analysis of neurophysiological data. Parametric methods were used for normal data, whereas we used the non-parametric Friedman test for non-normal data or small sample sizes. Post-hoc comparisons were tested with Wilcoxon tests. All tests were two-sided unless specified differently.

#### Theta power at rest and navigation

LFP segments of 2-second length during the resting period (inter-trial interval) and at trial onset were extracted for each trial. Power spectral densities were then estimated using the Welch’s method (*scipy*.*signal*.*welch* function) (6) with a Hanning window. The resulting estimate was smoothed (Gaussian kernel with sigma=0.3Hz), and the 2-10Hz frequency interval was averaged to assess the spectral density in the oscillatory band of interest (Fig 1-G).

#### Power spectral densities for low and high information densities

For each trial, the power spectral density was calculated (see above) for the initial 5 seconds of the signal. The obtained estimates were then split into 2 groups according to their information density condition (low: 5 or fewer cues/sec and high: more than 5 cues/sec). The group’s mean and S.E.M. for each frequency (2.6-9Hz) were computed and plotted in Fig 1-H.

#### Dominant theta frequency

To compute the dominant frequencies during virtual navigation, local maxima (peaks) along the theta range (2-10Hz) were identified using the *find_peaks* function from Scipy signal processing Python library. The dominant frequency was finally assessed by computing the center-of-mass (mean average) of the density peaks (Fig. 2A-F).

#### Time-varying instantaneous oscillation frequencies (frequency sliding)

Instantaneous frequency estimates were obtained following the methods in (7). The raw signal has been first bandpass filtered in the theta band. Next, the analytical signal is obtained by applying the Hilbert transform to the filtered data, and the phase angle time series is extracted. Frequency sliding is defined as the temporal derivative of the phase angle time series and converted to units of hertz by multiplying the derivative by the data sampling rate (in hertz) and dividing by 2pi. Finally, a median filter is applied to attenuate the noise spikes introduced by the frequency sliding which can appear as brief transient frequency jumps.

#### Bootstrap estimation analysis

All bootstrap-coupled estimation 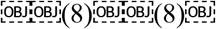. Specifi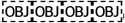cally, individua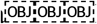l trials from each subject were grouped in the low (2 cues/sec) and high (14 cues/sec) information density and bootstrapped to assess the effect size of speed, number of visual cues, and information density (cue/sec) levels in modulating hippocampal theta dominant frequency (Fig 2D-F) and number of P-episodes (supplementary Fig S3).

For each subject, we derived the magnitude of the effect size (difference of the means between conditions) via the bias-corrected and accelerated bootstrap resampling technique, which creates multiple resamples from the set of observations (5000 repetitions), and computes the 95% CI for each of these resamples, where the effect size is presented as a bootstrapped 95% CI on a separate axis.

Further, 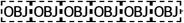population-level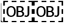 a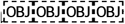nalyses u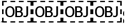sing the 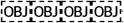same 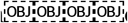meth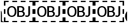od were p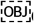erformed by aggregating the mean average of individual subjects at the lowest and highest condition level and estimating their respective effect size (Fig 2D-F and Fig S3, right-most column).

#### Gamma amplitude across the four theta quadrants

To compute the oscillatory amplitude of gamma (40-80Hz) in each of the four-cycle quadrants of theta (2-8Hz), we extracted the first 5-second LFP signal from each hippocampal electrode from each trial and notched filtered it (50Hz and respective harmonics) to remove power line interference.

Next, we applied the Hilbert transform onto the signal in both theta (2-8Hz) and gamma (40-80Hz) bands. With the Hilbert transform we obtain a complex time series:

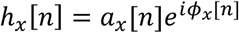

where *a*_*x*_[*n*]is the instantaneous analytic amplitude, and 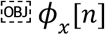 *ϕx*[*n*].

Theta phase was obtained by computing the instantaneous angle of the corresponding theta Hilbert signal in the complex space. Similarly, Gamma amplitude was computed by extracting the instantaneous magnitude of the gamma Hilbert signal.

For each trial, we then calculated the mean amplitude of gamma waves belonging to each of the four theta cycle quadrants (0-π/2, π/2-π, π-3π/2, 3π/2-2π). Mean gamma amplitudes per theta cycle quadrant of each subject were then grouped accordingly with the information density (cue/sec) condition (Fig. 3A).

#### Modulation index

The distribution of Gamma amplitudes along theta phase was computed using the *binned_statistic* function from the Scipy signal analysis library, set with a statistic method “mean” and 20 bins equally spaced along the cycle (0 to 2π).

The Modulation Index (MI) score was then calculated against a uniform distribution of the averaged gamma amplitude along the theta phase using the Kullback-Leibler divergence method (9) and used in (10, 11).

#### Oscillatory episodes analyses

Oscillatory power episodes in the gamma band (40-80Hz) were quantified using the P-episode methods previously described in (12–14). In summary, we extracted the LFP from each hippocampal electrode throughout the entire experimental session and notched filtered it (50Hz and respective harmonics) to remove power line interference. Next, we computed the time-frequency transform using Morlet wavelets defined by 20 oscillatory frequencies in the 40-80Hz range logarithmically spaced and with wavelet length set at half of its corresponding frequency. For each frequency, a 97th percentile amplitude threshold and a 3 cycles length of duration was set to identify oscillatory power events of marked and sustained activity (P-episodes). To do so, we percentiled the signal and set to zero all data points below the 97th amplitude threshold. Next, we convolved the resulting time-series of each frequency with their corresponding duration threshold (3 cycles) and identified P-episodes as the periods at which the signal survived zeroing. The number of P-episodes at each trial were then grouped (median) by the trial information density (cue/sec) level for each subject (Fig 3D).

#### Distribution of normalized P-episodes and theta frequencies

To assess the relationship between P-episodes and dominant theta frequency, we grouped and averaged (mean) the number of P-episodes present in each trial and the dominant theta from each subject at distinct information density (cue/sec) level. Next, we z-scored both the number of P-episodes and theta dominant frequency for each subject (Fig 3E-F).

#### Theta- P-episode coupling

Similarly to the theta-gamma coupling analysis (see Modulation index section), we assessed the instantaneous phase of hippocampal theta waves (2-8Hz) at the moment of each detected P-episode. For each 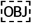trial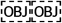, we computed the histogram distribution (count) of the number of P-episodes along the full oscillatory cycle (0-2π) (bins=21). The Theta- P-episodes coupling score was computed against a uniform distribution using the Kullb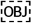ack-Leible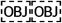r divergence method. Next, we grouped Theta- P-episodes coupling scores at the increasing levels of speed, number of visual cues, and information density (cue/sec) variables (Fig 3G).

### Computational model

The model used for the computational experiment is a spiking neural network consisting of a population of hippocampal pyramidal cells that receives global inhibitory feedback from two distinct populations of interneurons characterized by fast and slow dynamics, correspondingly (see Fig. 4a and Tables 2 and 3 below).

**Table 2.**
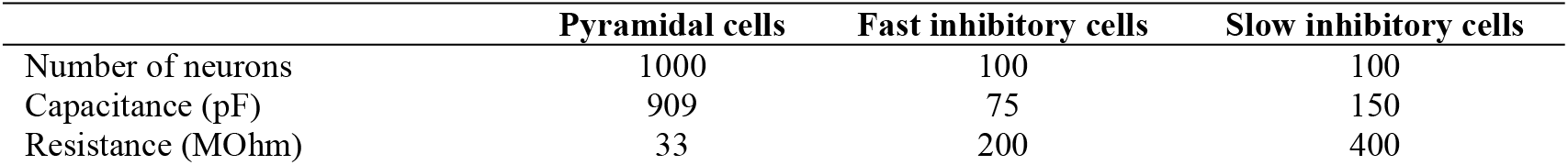
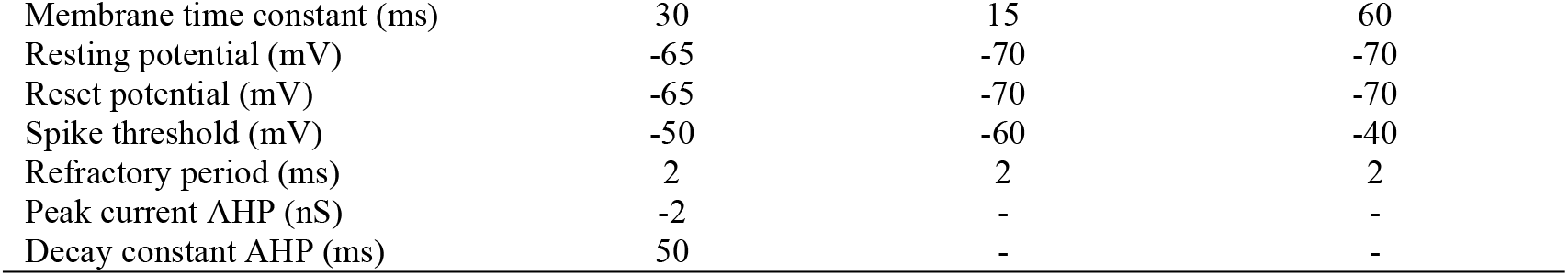
Parameters for the leaky integrate-and-fire neurons that compose each population. The parameters for the hippocampal pyramidal cells have been taken from previous literature. The parameters for the inhibitory interneurons were chosen so as to functionally reproduce fast and slow integration dynamics (15, 16) AHP = afterhyperpolarization.

|  | Pyramidal cells | Fast inhibitory cells | Slow inhibitory cells |
| --- | --- | --- | --- |
| Number of neurons | 1000 | 100 | 100 |
| Capacitance (pF) | 909 | 75 | 150 |
| Resistance (MOhm) | 33 | 200 | 400 |
| Membrane time constant (ms) | 30 | 15 | 60 |
| Resting potential (mV) | -65 | -70 | -70 |
| Reset potential (mV) | -65 | -70 | -70 |
| Spike threshold (mV) | -50 | -60 | -40 |
| Refractory period (ms) | 2 | 2 | 2 |
| Peak current AHP (nS) | -2 | - | - |
| Decay constant AHP (ms) | 50 | - | - |

**Table 3.** Parameters describing the synapses among the three populations. Synapses are modeled as exponentially decaying currents with instantaneous rise time (i.e., single exponential). The synaptic conductances reported here represent the maximum current values that the postsynaptic cells can receive at any point in time (drawn from a gaussian distribution with a coefficient of variation of 0.2). Therefore, individual synaptic efficacies are normalized by the number of presynaptic neurons. Every postsynaptic neuron receives connections from all the neurons of the corresponding presynaptic population. Parameters have been adapted from previous literature to reproduce the effects of fast and slow GABA*a* in the hippocampus (16). PSP = postsynaptic potential.

|  | E – Fast I | E – Slow I | Fast I – E | Slow I – E |
| --- | --- | --- | --- | --- |
| Type | AMPA | AMPA | GABA<sub>A</sub>,fast | GABA<sub>A</sub>,slow |
| Synaptic peak conductance (nS) | 4 | 2 | -4 | -3 |
| Decay time constant PSP (ms) | 3 | 10 | 3 | 50 |

Neurons are modeled as standard leaky integrate-and-fire units, characterized by the following differential equation:

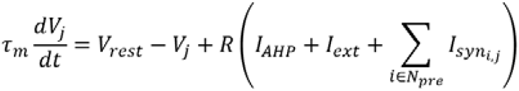

When the membrane voltage passes a threshold, it is reset to a specific value and it does not get updated during the following refractory period. Currents coming from presynaptic inputs are modeled as exponentially decaying potentials with instantaneous rise, as described by:

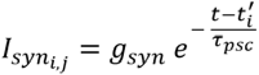

Where *t’* is the time of the presynaptic neuron’s last spike.

The stimulation protocol represents a person moving at a constant velocity through a linear track with overlapping place fields, where each item in the environment generates a place field of its own (as shown in Figure 4a). The skewness or asymmetry of the place fields (17) is approximated by a piecewise linear function (similar to (18)), with a maximum intensity of 2 nA at the peak. Each item consistently stimulates a random sample of 100 pyramidal cells from the pool of 1000 neurons. The stimulation is noisy, with an additive signal-dependent white noise with a coefficient of variation of 0.5. All items have the same exposure time (i.e., place field length) of 4 seconds. The total time needed to traverse the linear track is always 10 seconds (a trial). Hence, we set three different conditions where the information density (i.e., number of items) is varied (10, 20 and 30 items, representing the low, medium, and high conditions). Therefore, given the constant times of both the trials and the item exposures, increasing information density (i.e., adding more items) has the main effect of increasing the overlap between subsequent place fields.

In order to compare the behavior of the network across the different conditions, the compound potential (an LFP-like signal consisting of the average of all the membrane potentials) from the pyramidal population is extracted for further processing (see example in Fig. 4B), having 100 trials for each condition. Furthermore, frequency analyses are performed to the LFP-like signals to determine (1) the power spectral density distributions, (2) theta-gamma phase-amplitude coupling, (3) the center of mass of the power distributions in the theta range, and (3) the average number of gamma episodes per theta cycle. The range for theta is always set to 4-8 Hz, and for gamma is 30-80 Hz. The specific statistical methods are identical to the ones used to analyze the physiological data. Finally, the spike times are also extracted in order to correlate the theta phase of spikes against position within the place fields, testing for phase precession. All simulations were run using a customized spiking neural network simulator.

## References

1. Miller, G. A. The magical number seven, plus or minus two: Some limits on our capacity for processing information. Psychol. Rev. 63, 81 (1956).

2. Tulving, E. & Markowitsch, H. J. Episodic and declarative memory: role of the hippocampus. Hippocampus 8, 198–204 (1998).

3. Mormino, E. C. et al. Episodic memory loss is related to hippocampal-mediated β-amyloid deposition in elderly subjects. Brain 132, 1310–1323 (2008).

4. Taylor, M. J., Mills, T. & Pang, E. W. The development of face recognition; hippocampal and frontal lobe contributions determined with MEG. Brain Topogr. 24, 261 (2011).

5. Broadbent, N. J., Gaskin, S., Squire, L. R. & Clark, R. E. Object recognition memory and the rodent hippocampus. Learn. Mem. 17, 5–11 (2010).

6. O’Keefe, J., Burgess, N., Donnett, J. G., Jeffery, K. J. & Maguire, E. A. Place cells, navigational accuracy, and the human hippocampus. Philos. Trans. R. Soc. London. Ser. B Biol. Sci. 353, 1333–1340 (1998).

7. Mcnaughton, B. L., Battaglia, F. P., Jensen, O. & Moser, E. I. Path integration and the neural basis of the ‘ cognitive map ‘. Nat. Neurosci. 7, 663–678 (2006).

8. Pfeiffer, B. E. & Foster, D. J. Hippocampal place-cell sequences depict future paths to remembered goals. Nature 497, 74 (2013).

9. Johnson, A. & Redish, A. D. Neural ensembles in CA3 transiently encode paths forward of the animal at a decision point. J. Neurosci. 27, 12176–12189 (2007).

10. Eichenbaum, H. Time cells in the hippocampus: a new dimension for mapping memories. Nat. Rev. Neurosci. 15, 732 (2014).

11. Hoydal, O. A., Skytoen, E. R., Moser, M.-B. & Moser, E. I. Object-vector coding in the medial entorhinal cortex. bioRxiv 286286 (2018).

12. Danjo, T., Toyoizumi, T. & Fujisawa, S. Spatial representations of self and other in the hippocampus. Science (80-.). 359, 213–218 (2018).

13. Bos, J. J. et al. Multiplexing of Information about Self and Others in Hippocampal Ensembles. Cell Rep. 29, 3859–3871 (2019).

14. Alme, C. B. et al. Place cells in the hippocampus: eleven maps for eleven rooms. Proc. Natl. Acad. Sci. 111, 18428–18435 (2014).

15. Buzsáki, G. & Moser, E. I. Memory, navigation and theta rhythm in the hippocampal-entorhinal system. Nat. Neurosci. 16, 130 (2013).

16. Constantinescu, A. O., O’Reilly, J. X. & Behrens, T. E. J. Organizing conceptual knowledge in humans with a gridlike code. Science (80-.). 352, 1464–1468 (2016).

17. Aronov, D., Nevers, R. & Tank, D. W. Mapping of a non-spatial dimension by the hippocampal–entorhinal circuit. Nature 543, 719 (2017).

18. Rennó-Costa, C., Lisman, J. E. & Verschure, P. F. M. J. The mechanism of rate remapping in the dentate gyrus. Neuron 68, 1051–1058 (2010).

19. Lu, L. et al. Impaired hippocampal rate coding after lesions of the lateral entorhinal cortex. Nat. Neurosci. 16, 1085 (2013).

20. Bragin, A. et al. Gamma (40-100 Hz) oscillation in the hippocampus of the behaving rat. J. Neurosci. 15, 47–60 (1995).

21. O’Keefe, J. & Recce, M. L. Phase relationship between hippocampal place units and the EEG theta rhythm. Hippocampus 3, 317–330 (1993).

22. Lisman, J. & Buzsáki, G. A neural coding scheme formed by the combined function of gamma and theta oscillations. Schizophr. Bull. 34, 974–980 (2008).

23. Korotkova, T., Fuchs, E. C., Ponomarenko, A., von Engelhardt, J. & Monyer, H. NMDA receptor ablation on parvalbumin-positive interneurons impairs hippocampal synchrony, spatial representations, and working memory. Neuron 68, 557–569 (2010).

24. Robbe, D. & Buzsáki, G. Alteration of theta timescale dynamics of hippocampal place cells by a cannabinoid is associated with memory impairment. J. Neurosci. 29, 12597–12605 (2009).

25. Shirvalkar, P. R., Rapp, P. R. & Shapiro, M. L. Bidirectional changes to hippocampal theta–gamma comodulation predict memory for recent spatial episodes. Proc. Natl. Acad. Sci. 107, 7054–7059 (2010).

26. Pacheco Estefan, D. et al. Coordinated representational reinstatement in the human hippocampus and lateral temporal cortex during episodic memory retrieval. Nat. Commun. 10, 2255 (2019).

27. Wang, X.-J. & Buzsáki, G. Gamma oscillation by synaptic inhibition in a hippocampal interneuronal network model. J. Neurosci. 16, 6402–6413 (1996).

28. Buzsáki, G. & Wang, X. Mechanisms of gamma oscillations. Annu. Rev. Neurosci. 35, 203–25 (2012).

29. Buzsáki, G. Theta oscillations in the hippocampus. Neuron 33, 325–40 (2002).

30. Jeewajee, A., Barry, C., O’keefe, J. & Burgess, N. Grid cells and theta as oscillatory interference: electrophysiological data from freely moving rats. Hippocampus 18, 1175–1185 (2008).

31. McFarland, W. L., Teitelbaum, H. & Hedges, E. K. Relationship between hippocampal theta activity and running speed in the rat. J. Comp. Physiol. Psychol. 88, 324 (1975).

32. Bohbot, V. D., Copara, M. S., Gotman, J. & Ekstrom, A. D. Low-frequency theta oscillations in the human hippocampus during real-world and virtual navigation. Nat. Commun. 8, 14415 (2017).

33. Ekstrom, A. D. et al. Human hippocampal theta activity during virtual navigation. Hippocampus 15, 881–889 (2005).

34. Caplan, J. B. et al. Human theta oscillations related to sensorimotor integration and spatial learning. J. Neurosci. 23, 4726–36 (2003).

35. Aghajan, Z. M. et al. Theta oscillations in the human medial temporal lobe during real-world ambulatory movement. Curr. Biol. 27, 3743–3751 (2017).

36. Vass, L. K. et al. Oscillations go the distance: low-frequency human hippocampal oscillations code spatial distance in the absence of sensory cues during teleportation. Neuron 89, 1180–1186 (2016).

37. Monaco, J. D., Rao, G., Roth, E. D. & Knierim, J. J. Attentive scanning behavior drives one-trial potentiation of hippocampal place fields. Nat. Neurosci. 17, 725 (2014).

38. Watrous, A. J., Miller, J., Qasim, S. E., Fried, I. & Jacobs, J. Phase-tuned neuronal firing encodes human contextual representations for navigational goals. Elife 7, (2018).

39. Bush, D. et al. Human hippocampal theta power indicates movement onset and distance travelled. Proc. Natl. Acad. Sci. 114, 12297–12302 (2017).

40. Donoghue, T. et al. Parameterizing neural power spectra into periodic and aperiodic components. Nat. Neurosci. 2020 2312 23, 1655–1665 (2020).

41. Jutras, M. J., Fries, P. & Buffalo, E. a. Oscillatory activity in the monkey hippocampus during visual exploration and memory formation. Proc. Natl. Acad. Sci. 110, 13144–13149 (2013).

42. Cole, S. & Voytek, B. Cycle-by-cycle analysis of neural oscillations. J. Neurophysiol. 122, 849–861 (2019).

43. Tort, A. B. L. et al. Dynamic cross-frequency couplings of local field potential oscillations in rat striatum and hippocampus during performance of a T-maze task. Proc. Natl. Acad. Sci. U. S. A. 105, 20517–20522 (2008).

44. Tort, A. B. L., Komorowski, R. W., Manns, J. R., Kopell, N. J. & Eichenbaum, H. Thetagamma coupling increases during the learning of item-context associations. Proc. Natl. Acad. Sci. 106, 20942–20947 (2009).

45. Banks, M. I., White, J. A. & Pearce, R. A. Interactions between distinct GABA(A) circuits in hippocampus. Neuron 25, 449–457 (2000).

46. White, J. A., Banks, M. I., Pearce, R. A. & Kopell, N. J. Networks of interneurons with fast and slow γ-aminobutyric acid type A (GABA(A)) kinetics provide substrate for mixed gamma-theta rhythm. Proc. Natl. Acad. Sci. U. S. A. 97, 8128–8133 (2000).

47. Mehta, M. R., Quirk, M. C. & Wilson, M. A. Experience-dependent asymmetric shape of hippocampal receptive fields. Neuron 25, 707–715 (2000).

48. Lisman, J. E. & Jensen, O. The Theta-Gamma Neural Code. Neuron vol. 77 1002–1016 (2013).

49. Chadwick, A., Van Rossum, M. C. W. & Nolan, M. F. Flexible theta sequence compression mediated via phase precessing interneurons. Elife 5, (2016).

50. Sławińska, U. & Kasicki, S. The frequency of rat’s hippocampal theta rhythm is related to the speed of locomotion. Brain Res. 796, 327–331 (1998).

51. Bender, F. et al. Theta oscillations regulate the speed of locomotion via a hippocampus to lateral septum pathway. Nat. Commun. 6, 1–11 (2015).

52. Mann, E. O., Radcliffe, C. A. & Paulsen, O. Hippocampal gamma-frequency oscillations: from interneurones to pyramidal cells, and back. J. Physiol. 562, 55–63 (2005).

53. Tort, A. B. L., Rotstein, H. G., Dugladze, T., Gloveli, T. & Kopell, N. J. On the formation of gamma-coherent cell assemblies by oriens lacunosum-moleculare interneurons in the hippocampus. Proc. Natl. Acad. Sci. U. S. A. 104, 13490–13495 (2007).

54. Rotstein, H. G. et al. Slow and Fast Inhibition and an H-Current Interact to Create a Theta Rhythm in a Model of CA1 Interneuron Network. J. Neurophysiol. 94, 1509–1518 (2005).

## References

1. J. D. Monaco, G. Rao, E. D. Roth, J. J. Knierim, Attentive scanning behavior drives one-trial potentiation of hippocampal place fields. Nat. Neurosci. 17, 725 (2014).

2. A. Fedorov, et al., 3D Slicer as an image computing platform for the Quantitative Imaging Network. Magn. Reson. Imaging 30, 1323–1341 (2012).

3. X. D. Arsiwalla, et al., Network dynamics with BrainX3: a large-scale simulation of the human brain network with real-time interaction. Front. Neuroinform. 9, 02 (2015).

4. A. Principe, et al., Whole network, temporal and parietal lobe contributions to the earliest phases of language production. Cortex 95, 238–247 (2017).

5. A. Gramfort, et al., MEG and EEG data analysis with MNE-Python. Front. Neurosci. (2013) https://doi.org/10.3389/fnins.2013.00267.

6. P. Welch, The use of fast Fourier transform for the estimation of power spectra: a method based on time averaging over short, modified periodograms. IEEE Trans. Audio Electroacoust. 15, 70–73 (1967).

7. M. X. Cohen, Fluctuations in oscillation frequency control spike timing and coordinate neural networks. J. Neurosci. 34, 8988–8998 (2014).

8. J. Ho, T. Tumkaya, S. Aryal, H. Choi, A. Claridge-Chang, Moving beyond P values: data analysis with estimation graphics. Nat. Methods 16, 565–566 (2019).

9. S. Kullback, R. A. Leibler, On information and sufficiency. Ann. Math. Stat. 22, 79–86 (1951).

10. A. B. L. Tort, et al., Dynamic cross-frequency couplings of local field potential oscillations in rat striatum and hippocampus during performance of a T-maze task. Proc. Natl. Acad. Sci. U. S. A. 105, 20517–20522 (2008).

11. A. B. L. Tort, R. W. Komorowski, J. R. Manns, N. J. Kopell, H. Eichenbaum, Theta-gamma coupling increases during the learning of item-context associations. Proc. Natl. Acad. Sci. 106, 20942–20947 (2009).

12. L. K. Vass, et al., Oscillations Go the Distance: Low-Frequency Human Hippocampal Oscillations Code Spatial Distance in the Absence of Sensory Cues during Teleportation. Neuron 89, 1180–1186 (2016).

13. A. D. Ekstrom, et al., Human hippocampal theta activity during virtual navigation. Hippocampus 15, 881–889 (2005).

14. J. B. Caplan, et al., Human theta oscillations related to sensorimotor integration and spatial learning. J. Neurosci. 23, 4726–36 (2003).

15. C. Teeter, et al., Generalized leaky integrate-and-fire models classify multiple neuron types. Nat. Commun. 9, 1–15 (2018).

16. J. A. White, M. I. Banks, R. A. Pearce, N. J. Kopell, Networks of interneurons with fast and slow γ-aminobutyric acid type A (GABAA) kinetics provide substrate for mixed gamma-theta rhythm. Proc. Natl. Acad. Sci. 97, 8128–8133 (2000).

17. M. R. Mehta, M. C. Quirk, M. A. Wilson, Experience-dependent asymmetric shape of hippocampal receptive fields. Neuron 25, 707–715 (2000).

18. B. Melamed, S. Pan, Y. Wardi, HNS: A streamlined hybrid network simulator. ACM Trans. Model. Comput. Simul. 14, 251–277 (2004).

19. S. Cole, B. Voytek, Cycle-by-cycle analysis of neural oscillations. J. Neurophysiol. 122, 849–861 (2019).

